# Identification of signalling pathways involved in gill regeneration in zebrafish

**DOI:** 10.1101/2023.06.12.544629

**Authors:** Laura Cadiz, Maddison Reed, Simon Monis, Marie-Andrée Akimenko, Michael G. Jonz

## Abstract

The occurrence of regeneration of the organs involved in respiratory gas exchange amongst vertebrates is heterogeneous. In aquatic animals, such as amphibians and fishes, the gills regenerate completely following resection or amputation; whereas in mammals, only partial, facultative regeneration of lung tissue occurs following injury. Given the homology between gills and lungs, the capacity of gill regeneration in aquatic species is of major interest in determining the underlying molecular or signalling pathways involved in respiratory organ regeneration. In the present study, we used adult zebrafish (*Danio rerio*) to characterize signalling pathways involved in the early stages of gill regeneration. Regeneration of the gills, including the gas exchange surfaces, was induced by resection of the gill filaments and observed over a period of up to 10 days. We screened for the effects of the drugs, SU5402, dorsomorphin, and LY411575, which inhibit FGF, BMP or Notch signalling, respectively, on development of the blastema. Exposure to each drug for 5 days significantly reduced blastema formation in regenerates, compared to unresected controls. In separate experiments, and under normal conditions of regeneration, we used quantitative real-time PCR and observed an increased expression of genes encoding for the bone morphogenetic factor, Bmp2b, fibroblast growth factor, Fgf8a, a transcriptional regulator (Her6) involved in Notch signalling, and Sonic Hedgehog (Shha), in regenerating gills at 10 day post-resection, compared to unresected controls. *In situ* hybridization confirmed that all four genes were expressed in regenerating gill tissue. This study implicates BMP, FGF, Notch and Shh signalling in gill regeneration in zebrafish.

**SUMMARY STATEMENT:** Gill regeneration in zebrafish is mediated by multiple signalling pathways, including BMP, FGF, Notch and Sonic Hedgehog.

## INTRODUCTION

The fish gill is an organ that mediates multiple physiological processes, including exchange of O_2_ and CO_2_, control of respiration by peripheral chemoreceptors, and ion exchange (Evans et al., 2005). In teleost fish, the gill (or branchial) arches give rise to primary filaments, which in turn produce numerous secondary lamellae, where gas exchange occurs (Hughes, 1984). Although vastly different in morphology, the gill and mammalian lung are homologous structures that share similar cell types originating from endoderm-derived epithelium, and both develop from embryonic foregut (Gillis and Tidswell, 2017; Hockman et al., 2017; Kotton and Morrisey, 2014; Schittny, 2017). Unlike the lungs, which possess only facultative regenerative capacity (Reynolds et al., 2000; Beers and Morrisey, 2011; Kotton and Morrisey, 2014), the gills of adult fish are capable of complete regeneration of filaments and lamellae (Schäfer, 1936; Jonz et al., 2015; Stolper et al., 2019; Mierzwa et al., 2020; Nguyen and Jon, 2021). Given the evolutionary relationship between gills and lungs, investigation of the signalling pathways that orchestrate gill regeneration may enhance our understanding of the limitations imposed upon the regenerative potential of respiratory structures in other vertebrates.

Gill regeneration was first described for the external gills of the aquatic stage of *Necturus* (Eycleshymer, 1906). Regeneration of the internal gills in fish was later noted in goldfish (*Carassius auratus*; Schäfer, 1936) and more recently defined in the model vertebrates, zebrafish (*Danio rerio*; Jonz et al., 2015; Mierzwa et al., 2020; Nguyen and Jonz, 2021) and medaka (*Oryzias latipes*; Stolper et al., 2019). Gill regeneration follows the same general phases as described for other structures, which include wound healing, blastema formation and structural regrowth (Nye et al., 2003; Stocum, 2017; Cadiz and Jonz, 2020). In zebrafish, formation of a blastema, composed of undifferentiated or mitotic cells, begins within 24 h of gill filament resection (Jonz et al., 2015; Mierzwa et al., 2020). There is much evidence in the literature on wound healing in the fish gill following tissue damage or infection (e.g. Burkhardt-Holm et al., 1999; Dutta et al., 1996; Hemalatha and Banerjee, 1997), but replacement of differentiated cell types and regrowth of normal gill structure has only been described in zebrafish and medaka (Stolper et al., 2019; Mierzwa et al., 2020; Nguyen and Jonz, 2021). Nevertheless, little is currently known about the signalling mechanisms that drive complete regeneration in the gills of any aquatic species.

In the present study, we identified pathways involved in gill regeneration in zebrafish. We designed our approach based upon previous studies that have characterized pathways involved in regeneration, development or cell proliferation. We selected *fgf8a*, a gene that encodes a fibroblast growth factor (FGF) and is implicated in development of the branchial arches and head in zebrafish (Crump et al., 2004; Gebuijs et al., 2019). Two genes encoding bone morphogenetic protein (BMP), *bmp2b* and *bmp6*, were chosen for investigation based on evidence that they are involved in organogenesis (Chung et al., 2008) and fin regeneration (Smith et al., 2006), respectively. FGF and BMP also play a role in regeneration of the gills in axolotl (Saito et al., 2019). The genes, *jag1b* and *her6*, encoding a Notch ligand and effector, respectively, were selected because of their involvement in development of the branchial arches, or regeneration of the mandible, brain and caudal fin (Schebesta et al., 2006; Zuniga et al., 2010; Ueda et al., 2018; Kraus et al., 2022). Finally, a gene encoding Sonic Hedgehog signalling protein, *shha*, was investigated based on its role in regeneration of multiple tissues in zebrafish (Laforest et al., 1998; Armstrong et al., 2017; Dunaeva and Waltenberger, 2017; Thomas et al., 2018; Ueda et al., 2018; Iwasaki et al., 2018). Moreover, recent single-cell RNA-sequencing of the zebrafish gill revealed expression of genes encoding FGF, BMP, Notch, their receptors, as well as *shha* and *her6* (Pan et al., 2022). Using pharmacological inhibition of receptors implicated in regeneration, and selected gene expression analysis, we provide evidence for a role of *fgf8a*, *bmp2b*, *her6* and *shha* in gill regeneration in zebrafish.

## MATERIALS AND METHODS

### Animals

Adult wild-type zebrafish (*Danio rerio*) were bred and maintained in the Laboratory for the Physiology and Genetics of Aquatic Organisms at the University of Ottawa at 28.5°C on a 14:10 h light:dark cycle in a closed system. Zebrafish used in this study were 5–7 months of age. All animal handling and experimentation was approved by the University of Ottawa under protocol BL-3666 and performed in accordance with the Canadian Council on Animal Care.

### Gill resection and chemical exposure

Gill resection was performed following procedures described in Mierzwa et al. (2020). Briefly, zebrafish were lightly anesthetized using 0.1 mg ml^−1^ tricaine (MS 222; Syndel Laboratories Ltd., Qualicum Beach, BC) in system water. Zebrafish were then immobilized in a Petri dish in phosphate-buffered solution containing (in mM) 137 NaCl, 15.2 Na_2_HPO_4_, 2.7 KCl, 1.5 KH_2_PO_4_, balanced at pH 7.8 and with MS 222. Approximately 0.2–0.3 mm of tissue was resected from a single gill filament of the anterior hemibranch of the first branchial arch. Zebrafish were returned to 1-litre tanks for recovery and maintained for up to 10 days post-resection (d.p.r.) to allow for gill filament regeneration. Zebrafish were identified and monitored individually by skin pattern recognition.

In some experiments, the effects of pharmacological inhibitors on gill regeneration were tested. Chemicals were selected based on their selectivity for inhibiting known pathways involved in regeneration. Immediately after gill resection, groups of up to 5 zebrafish were placed in 1-litre tanks at 28.5°C and exposed to one of the following chemicals at concentrations tested in previous studies (sourced from MedChem Express LLC, Monmouth Junction, NJ, USA): 17 µmol l^−1^ SU5402 (cat. no. HY-10407) to inhibit FGFR1 receptor activation (Poss et al., 2000; Saera-Vila et al., 2016); 12 µmol l^−1^ dorsomorphin (DMD; cat. no. HY-13418A), an inhibitor of BMP type I receptors (Yu et al., 2008); or 5 µmol l^−1^ LY411575 (cat. no. HY-50752), a γ secretase inhibitor that blocks Notch signalling (Grotek et al., 2013). All drugs were first dissolved in dimethyl sulfoxide (DMSO) and added to system water. For controls, fish with resected gill filaments were maintained in water or 0.01% DMSO dissolved in water. Fish were exposed to drugs (or controls) until 5 d.p.r. with water changes every 24 h. No significant mortality was observed.

### Imaging and assessment of gill regeneration

The progress of blastema formation in control or drug treatment groups was assessed as a measure of gill regeneration. At 5 d.p.r., each individual was anesthetized and immobilized (as described above) a second time for live imaging. In this manner, gross morphological changes in filament structure were compared to filament structure immediately after resection. To enhance contrast of gill filaments during imaging, a small piece of tape coated with white nail polish was inserted between the two rows of filaments (hemibranchs) of the first branchial arch. Live imaging was performed using an AxioZoom.V16 with Apotome and Axiocam 506 (Zeiss, Jena, Germany).

Zebrafish were then prepared for measurement of the blastema in regenerating gill filaments. Animals were euthanized with 0.3 mg ml^−1^ tricaine in system water. The gill arch that had previously undergone filament resection was removed and washed with cold PBS. Tissue was fixed in 4% paraformaldehyde in PBS at 4°C overnight and then rinsed with PBS. Gill arches were mounted on glass slides in glycerol with a coverslip. The tips of gill filaments were examined using a Nikon A1 microscope (Nikon Instruments Inc., Tokyo, Japan).

Image J (Schneider et al., 2012) was used for image preparation and for measurement of blastema projection area. For each specimen, a region of interest was manually selected around the blastema and area was calculated using the Analyze-Measure tool. Statistical analysis was performed using Prism v.8.4.3 (GraphPad Software, San Diego, CA). Data sets were tested for normality using the D’Agostino and Pearson test. Significant differences between treatments were analyzed by one-way ANOVA, and Tukey’s test (P <0.05) was performed for *post-hoc* comparisons. Data are shown as means ± S.E.M.

### Reverse transcription quantitative polymerase chain reaction (RT-qPCR)

At 10 d.p.r., the first gill arch was removed from 8 zebrafish that had undergone the filament resection procedure. The first gill arch was also removed from 8 intact zebrafish for control. Gill arches were placed in cryotubes, frozen in liquid N_2_ and placed at –80°C until used. Total RNA from single gill arches was extracted using Trizol reagent (Thermo Fisher Scientific) and quantified using a NanoDrop 2000c UV-Vis Spectrophotometer (Thermo Fisher Scientific). cDNA was generated using QuantiTect Reverse Transcription Kit (Qiagen Inc., Toronto, ON, Canada) following manufacturers protocol. To check for genomic DNA contamination, a no-template negative control and a no reverse transcriptase negative control were included. mRNA gene expression of *fgf8a*, *bmp2b*, *bmp6*, *her6*, *jag1b* and *shha* in single gill arches was assessed by real time reverse transcription PCR (RT-qPCR) using SsoAdvanced Universal SYBR Green Supermix (Bio-Rad, Missisauga, ON, Canada). Expression of reference gene elongation factor 1a (*ef1a*) was stable between gills of control and regenerated groups and was therefore used to normalize mRNA expression for all genes. Standard curves were generated using a serial dilution of pooled cDNA to optimize primer reaction conditions. Each qPCR reaction included an initial step at 98°C to activate the enzymes in the mix, followed by 40 repeats of a denaturation step at 95°C and an annealing/extension step at the optimized temperature for each primer pair. A melt step from 66°C to 95°C in 0.5°C increments was included at the end of the reaction to check the specificity of the amplicon produced. Each biological sample was run in duplicate, and the relative abundance of mRNA was calculated using the ΔΔCt method (Livak and Schmittgen, 2001). See Table S1 for qPCR primer sequences. Statistical analysis was carried out using the Mann-Whitney U Test and Prism software.

### *In situ* hybridization

At 10 d.p.r., the first branchial arches were removed from zebrafish and prepared for whole-mount *in situ* hybridization using a protocol modified from Thisse and Thisse (2008). For each gene, gills from 3 zebrafish were sampled. Each gill arch was used for staining of regenerating filaments, as well as adjacent unresected filaments, which were used as intrinsic controls. Briefly, gills were fixed overnight in 4% paraformaldehyde at 4°C (Sigma, St. Louis, MO, USA) in diethyl pyrocarbonate in PBS (PBS-DEPC). The following day, gills were washed with PBS-DEPC, dehydrated in 100% methanol and stored at −20°C until used. Gills were then washed in PBS-DEPC and digested in 10 mg ml^−1^ proteinase K for 45 min at room temperature. Hybridization of probes was carried out at 67°C. After washes, samples were incubated at 4°C overnight with anti-digoxigenin (DIG) at 1:2,000 conjugated to alkaline phosphatase. Detection of hybridized RNA probes was achieved with with 225 μg ml^−1^ nitro blue tetrazolium (NBT) and 175 μg ml^−1^ 5-bromo 4-chloro 3-indolyl phosphate (BCIP) in Tris buffer. Staining was stopped when labelling was detected. Images were captured using an upright microscope (Axiophot, Zeiss, Jena, Germany) equipped with a DP-70 colour CCD camera (Olympus, Tokyo, Japan) and prepared using ImageJ.

Antisense RNA probes for *bmp2b* (1.3 kb; Laforest et al., 1998), *bmp6* (705 bp; Smith et al., 2006), *fgf8a* (1.5 kb; Zhang et al., 2010) and *shha* (2.5 kb; Smith et al., 2006) were synthesized, as previously described. A probe for *her6* (1.3 kb) was synthesized using a clone containing the complete coding sequence of *her6* (Dharmacon Clone ID: 4789975). The clone was linearized with BamHI and transcribed *in vitro* using T7 RNA polymerase. A 816 bp fragment of *jag1b* cDNA was amplified using *jag1b* forward 5’-CACGTGACGAGTTCTTTGGA-3’ and *jag1b* reverse 5’-CTGTGGCCATAGGTAAGTGG-3’ primers. This fragment was then used as a template to amplify a fragment including a T7 binding site at its end using the *jag1b-*L forward 5’-CACGTGACGAGTTCTTTGGACATTAT-3’ and *jag1b*-T7 reverse 5’-CATTATGCTGAGTGATATCCTGTGGCCATAGGTAAGTGGTTTAG-3’ primers. The *jag1b* probe was then transcribed *in vitro* using this template and T7 RNA polymerase.

## RESULTS

### Pharmacological treatment inhibited gill filament regeneration

Over the course of 5 days following the gill resection procedure, we observed normal regeneration of gill filaments, including the distal extension of the gill filament tip at the gross morphological level (Fig. 1A, B), and development of a blastema at higher magnification (Fig. 2A, B). Our observations were similar to those previously described for gill regeneration in zebrafish (Jonz et al., 2015; Mierzwa et al., 2020). For animals that were exposed to chemicals that targeted specific receptor types involved in regeneration or development, wound closure did not appear to be affected, but formation of the blastema was reduced at 5 d.p.r. Compared to zebrafish kept in system water or DMSO controls, zebrafish that were treated with SU5402, DMD or LY411575 displayed a negative effect of drug treatment upon the development of new tissue at the filament tip (Fig. 1C–E) and reduced blastema formation (Fig. 2C–E). At the concentrations tested, LY411575 had the greatest effect and produced a blastema with a mean area of 1,778 ± 268.5 µm^2^, significantly smaller compared to 8,081 ± 292.0 µm^2^ in DMSO controls (Fig. 2F; P<0.05; N=10). This was a 4.5-fold reduction in blastema area. SU5402 and DMD also caused the reduction of blastema area in regenerating filaments to 4,093 ± 371.4 µm^2^ and 3,479 ± 451.8 µm^2^, a 2.0-fold and 2.3-fold difference, respectively, compared to DMSO controls (Fig. 2F; P<0.05; N=10).

**Figure 1.**
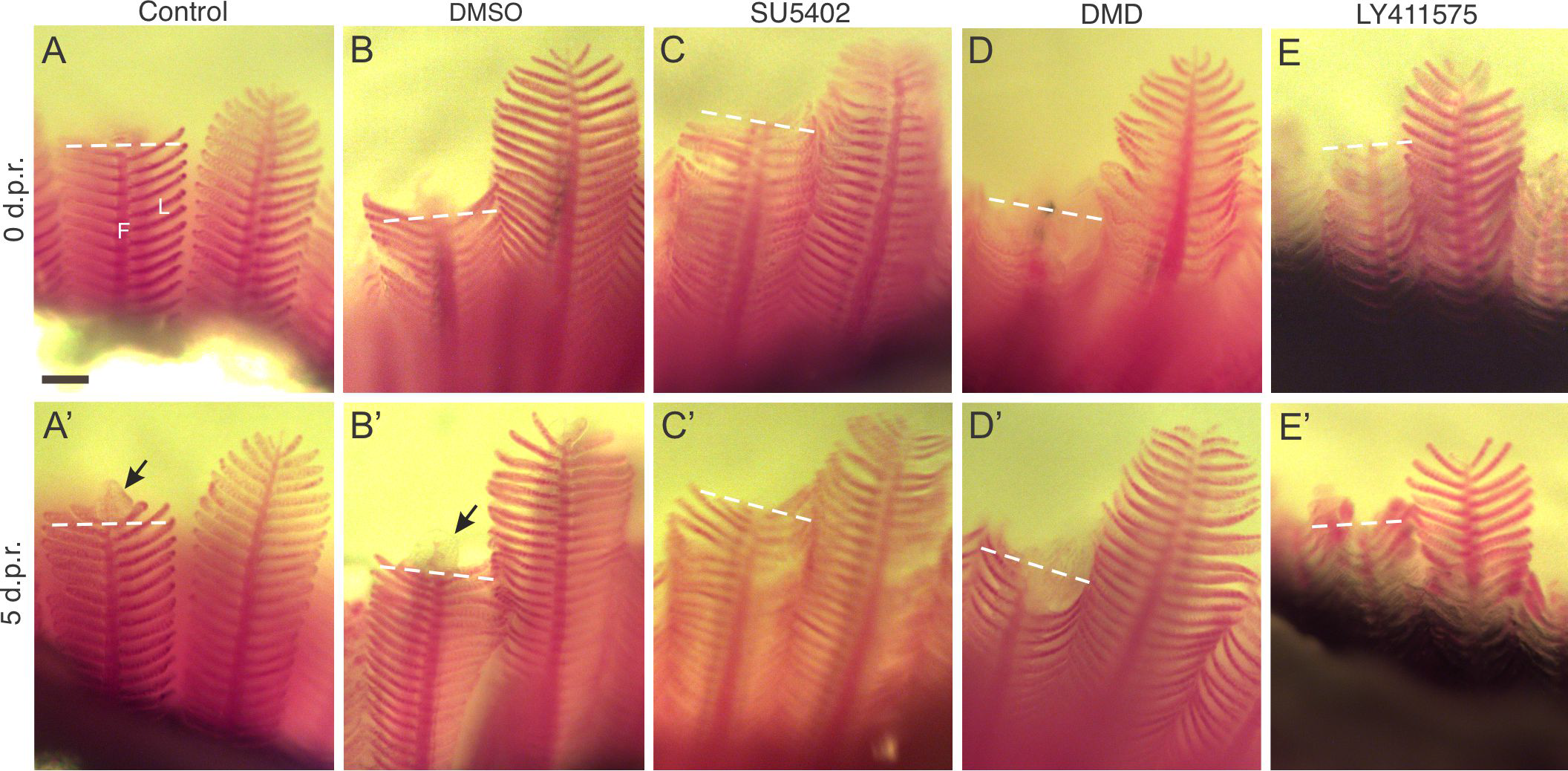
Pharmacological inhibitors restricted regeneration of the gill filaments. *In vivo* tracking of 5 individual zebrafish exposed to system water (A) or DMSO (B) as controls, or to SU5402 (C), dorsomorphin, DMD (D), or LY411575 (E). Gill filaments are shown immediately after the resection procedure (A–E) and again from the same individual at 5 days post-resection, d.p.r. (A’–E’). In all images, the dashed line indicates the site of resection. Development of a blastema (*arrows*) was observed only in controls (A’, B’) at this magnification. F, filament; L, lamella. Scale bar = 100 μm and applies to all panels.

**Figure 2.**
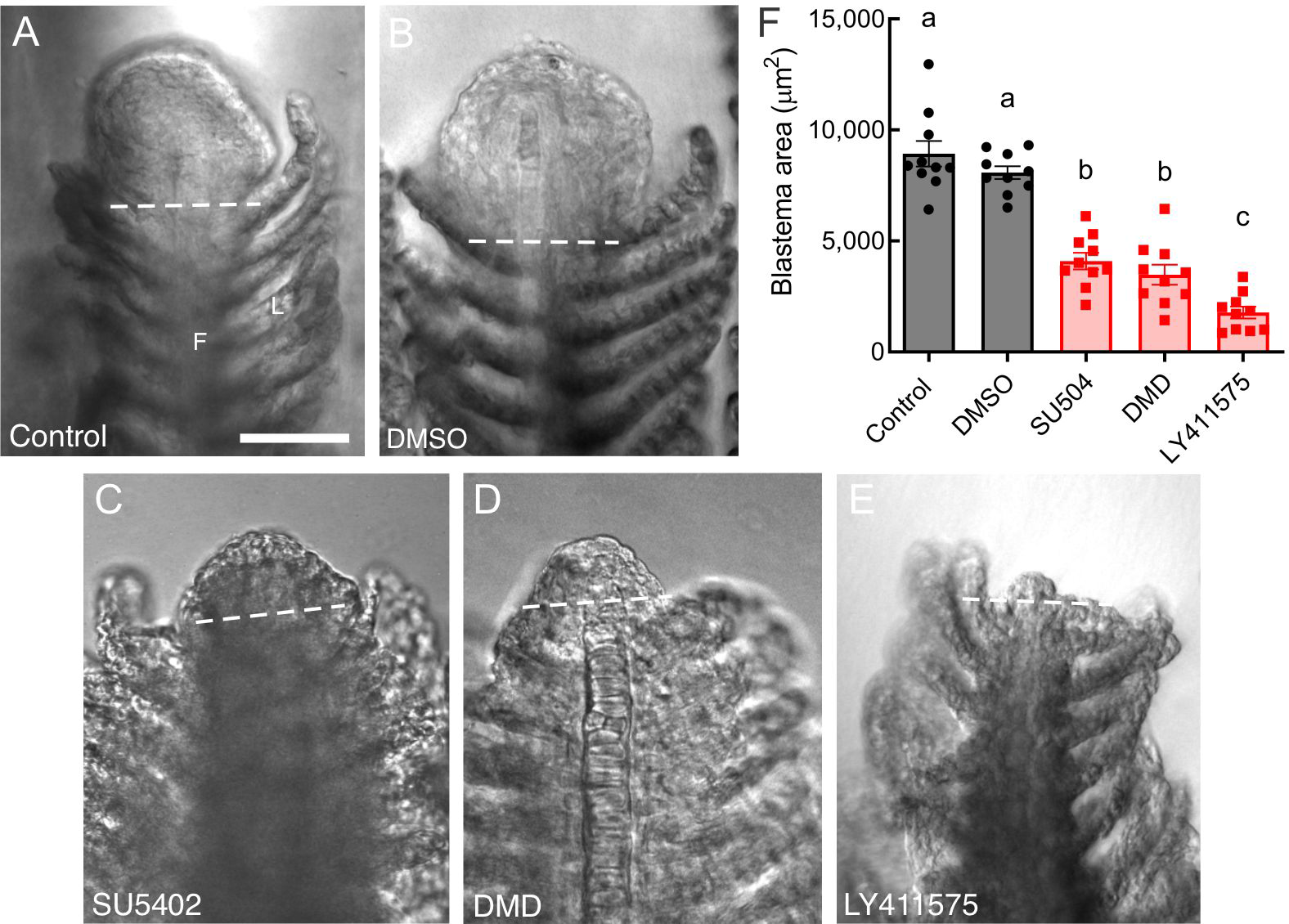
Pharmacological inhibitors reduced the area of blastema production during gill filament regeneration. The distal tips of gill filaments from zebrafish exposed to water (A) or DMSO (B) as controls, or to SU5402 (C), dorsomorphin, DMD (D), or LY411575 (E) are shown at 5 days post-resection (d.p.r.). A large blastema formed in controls (A, B) but was reduced in drug treatment groups. In all images, the dashed line indicates the site of resection. F, filament; L, lamella. Scale bar = 50 μm and applies to all panels. (F) Summary data showing mean ± S.E.M. blastema area (in μm^2^) at 5 d.p.r. for control and DMSO groups (*circles*), and for drug treatment groups (*squares*). Data were analyzed by one-way ANOVA and Tukey’s test (P<0.05; N=10 for each group). Significant differences between groups are indicated by different lower case letters.

### Gene expression in regenerating filaments

Based on results from chemical exposure experiments, which demonstrated reduced blastema formation by inhibition of FGF, BMP or Notch signalling, we further implicated these as important pathways involved in gill regeneration by quantifying relative expression of the genes, *fgf8a*, *bmp2b*, *bmp6*, *her6* and *jag1b* using RT-qPCR. *shha* was additionally included in this analysis because of the established role of this gene in regeneration. All six genes were expressed in the gills in both controls and regenerates (Fig. 3). In zebrafish that had undergone the resection procedure, the relative abundance of *fgf8a*, *bmp2b*, *her6* and *shha* significantly increased in the gills at 10 d.p.r., compared to unresected controls (Fig. 3A, B, D, F; Mann-Whitney U Test; P<0.05; N=8). Increases in abundance of these transcripts ranged from 4.3-fold for *bmp2b* to 10.9-fold for *her6*.

**Figure 3.**
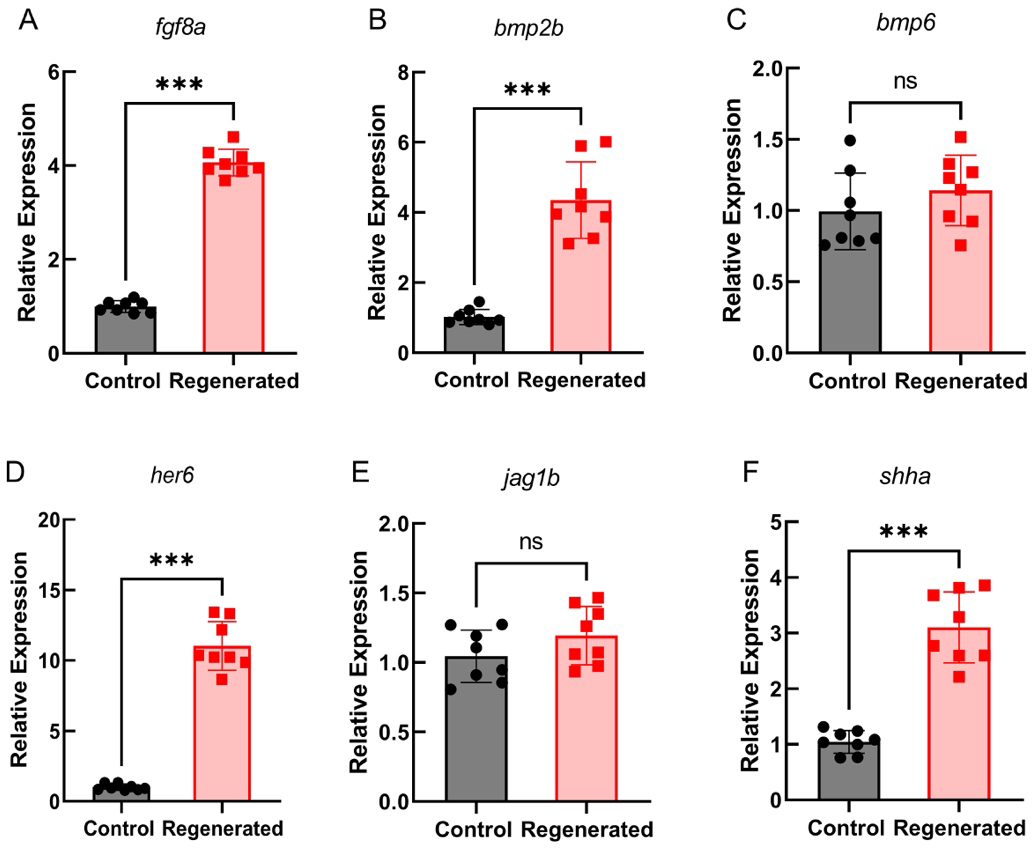
Gene expression analysis. Relative mRNA expression of (A–F) *fgf8a*, *bmp2b*, *bmp6*, *her6*, *jag1b* and *shha* in regenerating gills of zebrafish 10 days after the resection procedure (*squares*), compared to expression in gills from intact animals (*circles*). Expression of all genes was observed under control conditions, but relative abundance increased during regeneration for only *fgf8a*, *bmp2b*, *her6* and *shha*. Data were normalized to the mRNA abundance of the reference gene, *ef1a*. Data were analyzed using the Mann-Whitney U Test and means ± S.E.M. significantly different from control are indicated by asterisks (P<0.01; N=8 for each group). ns = not significant.

Our next goal was to identify domains of gene expression in the gills using *in situ* hybridization (Fig. 4). With the exception of *shha*, expression of other genes was below the level of detection in the distal region of gill filaments in unresected animals, but we observed clear expression of *fgf8a*, *bmp2b* and *her6* near the site of resection in gill filaments that were undergoing regeneration (Fig. 4A, B, D). The expression domains of these genes corresponded to the site of blastema formation at the distal tips of the filaments. Intense labelling of *shha* was also found in regenerating filament tips, as well as along the filament epithelium proximal to the site of resection (Fig. 4F). Weak labelling was also found along the filament epithelium in unresected controls (Fig. 4F’), suggesting that *shha* expression was not limited to regenerating tissue in the gill.

**Figure 4.**
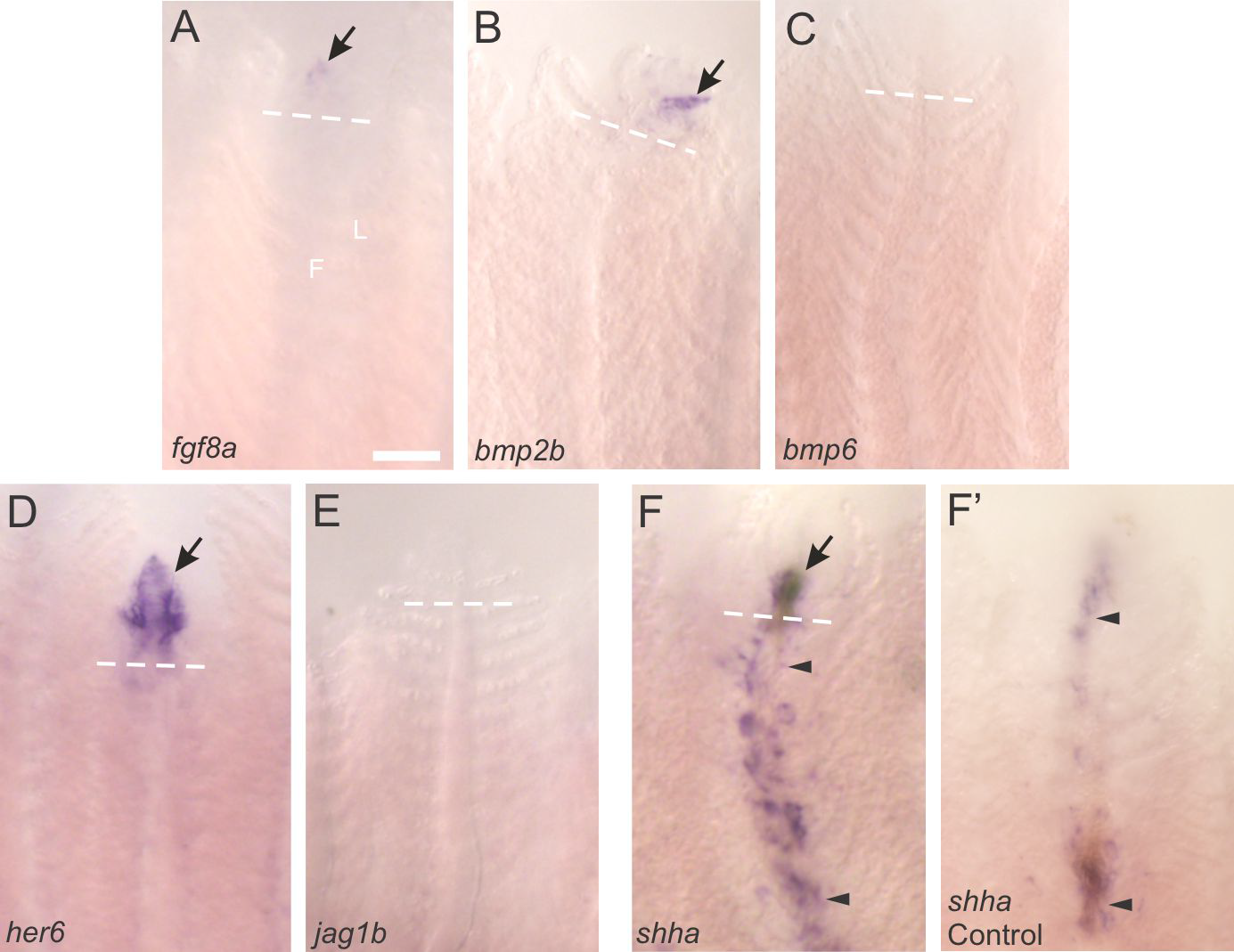
Expression domains in regenerating gill filaments as shown by *in situ* hybridization. The distal tips of previously-resected gill filaments at 10 days post-resection (d.p.r.) were stained for expression of (A–F) *fgf8a*, *bmp2b*, *bmp6*, *her6*, *jag1b* and *shha*. Dashed lines indicate the site of resection. Gene expression (*arrows*) was detected in the regenerating filament tips in panels A, B, D and F. For *shha*, expression was observed along the length of the filaments (*region between arrowheads*) in regenerating filaments (F) and in unresected controls (F’), in addition to the increased expression in regenerating tissue (F, *arrow*). F, filament; L, lamella. Scale bar = 100 μm and applies to all panels.

## DISCUSSION

The present study used chemical exposures and gene expression analysis to identify signalling pathways involved in gill regeneration in zebrafish. While all genes tested were expressed in the gills, we demonstrated an increase in the relative abundance of four genes— *fgf8a*, *bmp2b*, *her6* and *shha*—in regenerating filaments and provide evidence for a role for FGF, BMP, Notch and Shh signalling in gill regeneration.

Development and growth of the gills in fish is well described in the literature (e.g. Morgan, 1974; Rombough, 1988; Kimmel et al., 1995; Evans et al., 2005; Jonz and Nurse, 2005; Fu et al., 2010; Hwang and Chou, 2013; Mongera et al., 2013; Kwong and Perry, 2015; Gillis and Tidswell, 2017; DeLaurier, 2018; Stolper et al., 2019; Lelièvre et al., 2023). By contrast, the regenerative potential of the gill is just beginning to be described at the cellular and molecular level. In zebrafish, gill regeneration begins immediately after gill filament resection or amputation and results in formation of a blastema at the filament tip within 24 h. Approximately half of resected gill tissue is replaced by 40 d.p.r., and about 85% is replaced by 160 d.p.r. (Mierzwa et al., 2020). Regeneration includes a return to normal gill morphology, including vascularization, perfusion, innervation, and the return of multiple cell types, such as chemoreceptors, neurons and ionocytes (Mierzwa et al., 2020; Nguyen and Jonz, 2021).

In the present study, we used a pharmacological approach to identify signalling pathways potentially involved in mediating gill regeneration. Each drug partially reduced or delayed blastema formation, indicating that multiple pathways are likely involved in the regenerative process. SU5402 inhibits enzymatic activity of receptor tyrosine kinases, including FGFR1 receptors, and has been shown to impair blastema formation, and fin and muscle regeneration in zebrafish (Poss et al., 2000; Saera-Vila et al., 2016). In the gills, we found that SU5402 reduced blastema production to about half its normal size. Multiple types of FGF receptors, including *fgfr1*, *fgfr2*, *fgfr3* and *fgfr4*, are expressed in the zebrafish gill and were localized to cell types, such as fibroblasts, endothelial cells and pavement cells (Pan et al., 2022). These receptors may represent potential targets of SU5402 activity. Our demonstration of *fgf8a* expression within the developing blastema suggests that Fgf8 may be an endogenous regulator of gill regeneration at the site of resection. Fgf8 has been shown to be involved in development of the branchial arches and head in zebrafish (Crump et al., 2004; Gebuijs et al., 2019), and gill regeneration in axolotl (Saito et al., 2019).

Dorsomorphin is a highly-selective inhibitor of BMP type I receptors that perturbs embryogenesis in zebrafish when applied *in vivo* (Yu et al., 2008). In addition, the gene encoding its ligand, *bmp2b*, is required for organogenesis in zebrafish (Chung et al., 2008) and has already been implicated in gill regeneration in axolotl (Saito et al., 2019). The present study demonstrated that DMD significantly reduced blastema progression in the gill, and we localized expression of *bmp2b* specifically to the site of regeneration in gill filaments. Given the recent report of expression of *acvrl1*, a gene encoding a subtype of BMP type I receptors, in the zebrafish gill (Pan et al., 2022), the BMP pathway represents a likely candidate for regulation of gill regeneration in zebrafish.

We used LY411575 as a γ secretase inhibitor to block the Notch signalling cascade. This drug was used in a previous study to demonstrate that Notch signalling is required for the proliferation of blastema cells during fin regeneration, where they also found robust expression of *her6*—a gene encoding a Notch effector protein—confined to the proliferative zone of the blastema (Grotek et al., 2013). *her6* has also been implicated in neurogenesis and bone formation during mandibular regeneration in zebrafish (Ueda et al., 2018; Kraus et al., 2022). In agreement with Grotek et al. (2013), we found that LY411575 almost entirely inhibited growth of the blastema in the gill. Consistent with the dramatic effects of Notch inhibition, under normal conditions of regeneration we report a 10.9-fold increase in relative abundance of *her6* in regenerates compared to the intact gill, and intense expression of this gene that was confined to the blastema at 10 d.p.r. Interestingly, using single-cell RNA-sequencing, *her6* expression was shown in cells of the zebrafish gill, such as endothelial cells and fibroblasts, and *her6* expression increased in the gills following *in vivo* exposure to hypoxia (Pan et al., 2022), a stimulus known to induce cell proliferation or growth in gill cells (Jonz et al., 2004; Regan et al., 2011; Porteus et al., 2014). Together, these data argue strongly for a role for Notch signalling in gill regeneration by mediating blastema cell proliferation.

Investigation of potential Shh signalling was included in the present study because of its important role in regeneration of multiple tissues in zebrafish, and its role in cell proliferation and neurogenesis (Laforest et al., 1998; Armstrong et al., 2017; Dunaeva and Waltenberger, 2017; Thomas et al., 2018; Ueda et al., 2018; Iwasaki et al., 2018). From our *in situ* hybridization experiments, we show that *shha* staining increased within the regenerating filament tip at 10 d.p.r., and this was in line with our gene expression analysis that indicated an increase in the relative abundance of *shha* in regenerates. However, we also observed weak *shha* staining along the length of the gill filaments in both unresected controls and regenerates. This suggests that, while *shha* may participate in gill regeneration at the filament tip, it must also play another role elsewhere in the gill. Interestingly, RNA-sequencing data indicated that *shha* was expressed only in chemoreceptive neuroepithelial cells (Pan et al., 2022). Neuroepithelial cells in the gills are found along the length of the gill filament and at the tip, and function as sensors of hypoxia (Jonz et al., 2003; Jonz et al., 2004). As part of their role in initiating acclimatization to low environmental oxygen, gill neuroepithelial cells proliferate and undergo hypertrophy when confronted with chronic hypoxia (Jonz et al., 2004; Pan et al., 2021). Given the role of Shh in mediating cell proliferation, future studies may seek to identify whether Shh may be important in regulating the population of oxygen chemoreceptors in the gills. Another possibility is that expression of *shha* along the length of the gill filament represents regions of constitutive cell proliferation to maintain multiple cell populations. Stolper et al. (2019) identified homeostatic stem cells along the gill filaments in medaka, and similar mitotic cells were labelled with antibodies against the proliferating cell nuclear antigen in zebrafish (Mierzwa et al., 2020).

We have demonstrated that multiple pathways are involved in gill regeneration in zebrafish, and have presented evidence that identifies a role for FGF, BMP, Notch and Shh signalling in mediating this process. The molecular basis of gill regeneration in fish is still in its early stages. Continued studies on gill regeneration will improve our understanding of cell proliferation and tissue replacement in the gills following injury or disease in fish. Moreover, many of the genetic pathways that promote gill and lung regeneration appear to be highly conserved in vertebrates. For example, in mammalian lung, BMP signalling is critical for stem cell activation and differentiation following damage (Chung et al., 2018), whereas FGF promotes tissue repair in the lung after injury (Finch et al., 2013). In addition, Notch controls epithelial cell transdifferentiation in the injured lung (Kiyokawa and Morimoto, 2020). Given the regenerative capacity of the zebrafish gill, and the genetic tools available for this model vertebrate, future research on gill regeneration in zebrafish may lead to a better understanding of the relatively limited regenerative potential of the lung, which may lead to new developments in treating human lung disease.

### Author contributions

Conceptualization: M.G.J., L.C.; Methodology: all authors; Validation: L.C., M.R.; Formal analysis: L.C., M.R.; Investigation: all authors; Resources: M.G.J., M.A.A.; Data curation: L.C., M.R.; Writing - original draft: M.G.J., L.C., M.R.; Writing - review & editing: all authors; Supervision: M.G.J., M.A.A.; Project administration: M.G.J.; Funding acquisition: M.G.J.

## Data availability

All data associated with this work are presented in the manuscript.

## Supplemental

**Table S1.**
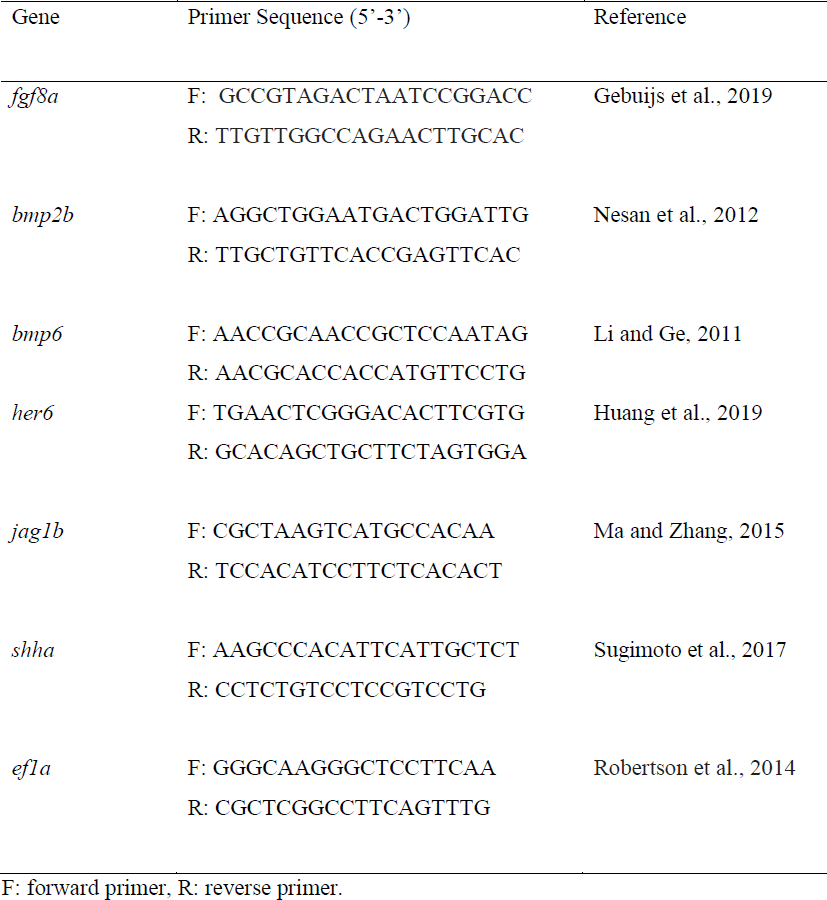
Primer pairs used for mRNA quantification by RT-qPCR.

